# Fenbendazole resistance in *Heterakis gallinarum*, the vector of *Histomonas meleagridis*, the causative agent of Blackhead Disease in poultry

**DOI:** 10.1101/2021.03.30.437780

**Authors:** James B. Collins, Brian Jordan, Andrew Bishop, Ray M. Kaplan

## Abstract

Due to their ubiquity, management of parasites is a common and important factor for profitable production of poultry. *Heterakis gallinarum*, the cecal nematode, is the most common nematode parasite of poultry. While typically causing no pathology on its own, *H. gallinarum* is the vector of *Histomonas meleagridis*, a protozoan parasite that causes blackhead disease. *Histomonas meleagridis* is highly pathogenic in turkeys, potentially causing high mortality. In contrast, disease caused by *H. meleagridis* is much less severe in chickens, where it primarily reduces productivity without manifestations of clinical disease. There are no approved treatments for *H. meleagridis*, making control reliant on control of the helminth vector through the use of fenbendazole (FBZ) the only drug labeled for treatment of *H. gallinarum* in the United States We were contacted by an industry veterinarian regarding health-related concerns in a broiler-breeder house due to histomoniasis, despite frequent anthelmintic treatments. Since we had recently diagnosed resistance to FBZ in *Ascaridia dissimilis*, a closely related nematode of turkeys, we were interested to determine if *H. gallinarum* had also evolved resistance to FBZ. *Heterakis gallinarum* eggs were isolated from litter collected from the house and used to infect 108 chickens. Treatment groups included a non-treated control, a label-dose and a 2X-label dose of FBZ, with 36 birds per group divided into two replicate pens of 18 birds each. Birds were placed at 1-day post hatch, and at 3 weeks of age were infected with 150 embryonated eggs via oral gavage. Two weeks post infection, treated birds were administered a minimum of either a label- or 2X label-dose of FBZ in water for 5 days (SafeGuard^®^ Aquasol, 1mg/kg BW). To ensure that all birds consumed the full intended dose at a minimum, the dosage was calculated using 1.25 times the average body weight. One-week post treatment, birds were euthanized, ceca removed, and parasites enumerated. Efficacy was calculated by comparing the total numbers of worms recovered from each treatment group to the numbers recovered in the non-treated control group. There were no significant differences in worm numbers recovered from any of the three groups (p-value=0.81). There also was no efficacy benefit to treatment with a 2X dose; *H. gallinarum* worm counts were reduced by 42.7% and 41.4%, for the label and 2X dosages, respectively. These data provide strong evidence that *H. gallinarum* has developed resistance to FBZ. Consequently, in houses infected with FBZ-resistant *H. gallinarum*, *H. meleagridis* will be able to cycle through the birds in an unrestricted manner. Further investigation is needed to determine the prevalence of resistance in *H. gallinarum on* chicken farms, but it is clear this has the potential to have a large-scale economic impact on the poultry industry. These data when viewed together with our recent findings of FBZ resistance in *A. dissimilis*, suggest that drug resistance in ascarid nematodes may be an important emerging problem on poultry operations.

## 1. INTRODUCTION

The near ubiquity of parasites on poultry farms makes the management of parasites a common and important factor effecting profitable production of poultry. A study of birds from 10 different production companies Southeastern United Stated reported that 98.6% of birds are infected by parasitic helminths, with 96% being infected with the cecal worm, *Heterakis gallinarum* (Yazwinski, et al., 2013). *Heterakis gallinarum* belongs to the Family Ascarididae, which also contains the closely related *Ascaridia galli* and *Ascaridia dissimilis*, important small intestinal nematodes of chickens and turkeys, respectively. Helminth eggs from this family are resistant to environmental pressures such as temperature, dehydration, and pH extremes, causing a cycle of continuous infection and transmission within the house environment (Cauthen, 1931; Tarbiat, et al., 2015). *Heterakis gallinarum* is a small nematode that rarely causes significant direct pathology, but it serves as the vector for *Histomonas meleagridis*, a highly pathogenic protozoal parasite that is the causative agent of Blackhead disease in poultry.

*Histomonas meleagridis* currently ranks as the highest research priority in broilers of any parasite of poultry (Armour, et al., 2020). *Histomonas meleagridis* is carried within the eggs of *H. gallinarum*, and Histomonads are released into the gut when the larvae hatch from the nematode egg in the intestine. *Histomonas meleagridis* causes the disease histomoniasis, which is characterized by necrosis in the mucosal tissues of the ceca and liver, and may cause dark discoloration of the head, hence the name Blackhead. Historically, infections in turkeys often produced high levels of mortality, whereas in chickens, infection was largely asymptomatic. Recently, this view has shifted as studies show that both chickens and turkeys demonstrate clinical signs such as apathy, depression, and ruffled feathers, together with decreased feed and water uptake (Liebhart, et al., 2017). *Histomonas meleagridis* has now been shown to impact chickens in multiple different production systems, including reduced feed conversion in broilers, and decreased egg quality and production in layers and breeders, both caged and cage-free (Clark and Kimminau, 2017; Grafl, et al., 2011; Liebhart, et al., 2013) Despite these significant effects on health and production, there are currently no FDA approved treatments for histomoniasis, making control of this diseases dependent on control of the *H. gallinarum* vector.

Currently, fenbendazole (FBZ) is the only anthelmintic approved for use against ascarids of poultry in the United States. Fenbendazole belongs to the benzimidazole class of anthelmintics, a drug class used widely across multiple livestock species. In registration studies for SafeGuard^®^ Aquasol^®^, a formulation of FBZ that is suspended in water for delivery, average efficacy against *H. gallinarum* was 96.2%, similar to that of *Ascaridia galli*, at 97.6% (United States Food and Drug Administration, 2018). Likewise, in registration studies for the feed additive formulation of SafeGuard^®^, average efficacy against *H. gallinarum* was 97.85% in growing turkeys (United States Food and Drug Administration, 2000). Both formulations are delivered from a central ration, or medication tank, and then distributed throughout the house. These methods of administration, along with human error, may result in poor delivery of treatment, leading to underdosing. Underdosing is recognized as one of the major contributors to the development of anthelmintic resistance (Jackson and Coop, 2000; Silvestre, et al., 2001), and combined with the high frequency of treatment, in as little as every four weeks, development of resistance is a major concern (Smith, et al., 1999).

Resistance to benzimidazoles in many of the most economically important strongylid nematodes of livestock is highly prevalent (Howell, et al., 2008; Kaplan, 2004; Kaplan and Vidyashankar, 2012), however resistance in ascarid nematodes appears to be rare. In poultry, reduced efficacy was first reported in the turkey nematode *Ascaridia dissimilis*, leading to speculation that drug resistance may have developed (Yazwinski, Tucker et al. 2013). This suspicion was recently confirmed in a controlled efficacy study, where FBZ resistance was clearly demonstrated in *A. dissimilis* (Collins, et al., 2019). This confirmation of resistance highlights the potential of ascarid nematodes of poultry to develop resistance to FBZ, and since birds treated with FBZ may be infected with both *A. dissimilis* and *H. gallinarum*, FBZ resistance in *H. gallinarum* may already exist.

Given the recent concern of increased infection and disease from *H. meleagridis* in breeder chickens and having demonstrated FBZ resistance in one ascarid species of poultry, we wanted to determine if *H. gallinarum*, had also developed resistance. Through collaboration with industry veterinarians, we identified a farm with suspected-resistant *H. gallinarum* and conducted a controlled efficacy trial to determine if the worms on that farm were in fact resistant to FBZ.

## 2. MATERIALS AND METHODS

### 2.1 Chickens

One hundred eighteen, Cobb 500, chicks were hatched and placed the following day in housing at the Poultry Science Farm at the University of Georgia. Nipple drinkers and hanging feeders were used to provide water and feed *ad libitum*. Birds were fed a diet of non-medicated Nutrena^®^ NatureWise^®^ Chick Starter Grower feed.

### 2.2 Parasite Isolates

A potentially resistant isolate of *Heterakis gallinarum*, AmFa 1.0, was identified through collaboration with an industry veterinarian. Prior to May of 2017, the farm of origin for this isolate, treated birds with a variety of treatments including FBZ, but after May of 2017, exclusively treated six flocks, four treatments per flock, with FBZ. Litter was obtained from the suspect farm, and eggs were isolated using previously established protocols (Collins et al., 2019). Briefly, litter was washed through a series of sieves to remove debris, and then the remaining sediment was added to a solution with specific gravity of 1.15 and centrifuged at 433g for 7 mins. Eggs within the fluid phase were collected on a 32uM sieve, rinsed with deionized water, and stored in tissue culture flasks in deionized water containing 0.5% formalin at 10°C. Prior to infecting the birds, flasks were incubated at 30°C for four weeks, until eggs were fully developed, and then stored at 10 °C until infection.

### 2.3 Infection & Treatment

Birds were divided into two replicates of 18 birds each for the following three treatments: non-treated control, label dosage of FBZ, and 2x label dosage of FBZ. An additional 10 birds remained uninfected as environmental controls to confirm that there was no prior contamination of the study environment with *H. gallinarum* eggs. Birds were allowed to grow to three weeks of age before being infected with approximately 150 embryonated eggs via oral gavage. Mesh curtains were placed between pens to prevent any cross-over of birds between treatment groups.

Two weeks post infection, birds were treated with either the label or a 2x dose of SafeGuard^®^ Aquasol^®^ (Label Dosage: 1mg/kg BW), a FBZ formulation designed for delivery in water. To increase the likelihood that every bird received the target dosage at a minimum, the average bird weight of the groups on the day before treatment plus 25% was used to calculate the dose administered. Calculated volumes of FBZ were mixed into 90% of the volume of water estimated to be consumed, as per production guidelines. For delivery of the FBZ, water lines were connected to carboys with both replicates of each treated group receiving water from the same carboy. As per label directions, treatment was administered over the course of five days, resuspending the drug daily.

### 2.4 Worm Recovery

Seven days post-treatment, all birds were humanely euthanized for worm recovery. Ceca were removed, opened, and placed in physiological saline. Samples were incubated overnight at 37C to aid in the recovery of tissue associated nematodes. Cecal contents were then washed over a 50uM mesh sieve to remove small debris, and cecal cores if present were manually disrupted. Contents were then examined under a dissecting microscope, and all nematodes were recovered and enumerated.

### 2.5 Statistical Analysis

Differences between the three groups were evaluated using Kruskal-Wallis test by ranks with Dunn’s correction, a non-parametric analysis of variance with multi-comparisons (Graph Pad Prism 8, San Diego,CA). Analysis of control vs. label dosage and control vs. 2x dosage were done with the following parameter: Two-samples unpaired with zero-inflation, and a correction factor of 1. Efficacy was calculated using eggCounts, an R package using a Bayesian hierarchical model to determine anthelmintic efficacy (Wang, et al., 2018).

## 2. RESULTS

No significant differences were observed between the three groups (p>0.81). Model adjusted efficacies for the label dose and 2x label dose groups were 42.7% and 41.4%, with upper 95% CI of 74.2 and 74.1%, respectively (Table 1). Both of these upper 95% CIs are well below reported efficacies of 96.2% and 97.85%, for the water and feed formulations, respectively (United States Food and Drug Administration, 2000; United States Food and Drug Administration, 2018). These data provide strong evidence that these *H. gallinarum* are resistant to FBZ.

**Table 1.** Zero-inflated adjusted average worm count and efficacy.

| Treatment | Mean Number of<br>Worms | % Reduction | 95%CI |
| --- | --- | --- | --- |
| Control | 12.2 | - | - |
| Label Dosage | 6.8 | 42.7% | (4.5%,74.2%) |
| 2x Dosage | 6.8 | 41.4% | (3.4%,74.1%) |

## 4. DISCUSSION

We confirm here for the first time, resistance to FBZ in the poultry nematode, *Heterakis gallinarum*. Efficacies of the label dosage and 2X the label dosage were 42.7% and 41.4% respectively, with no significant difference between the two dose levels, nor between these two treated groups and the non-treated controls (p>.9999). The lack of differentiation in efficacy for the two dose levels indicate not only that resistance has developed, but there appears to be a virtual total lack of efficacy. The observed efficacies are likely a due to random variation between birds and groups, as the worm counts were highly over-dispersed with many zeros in all three groups (Supplementary table 1). Similar to the resistant isolate of *A. dissimilis* we previously identified, this farm of origin has a history of FBZ use, further highlighting the risks for resistance associated with having only one approved compound for use against helminths of poultry. As compared to *A. dissimilis, H. gallinarum*, by itself, poses less disease risk to its host. However, due to its role as a vector for *H. meleagridis*, FBZ resistance in *H. gallinarum* poses important health challenges in poultry operations.

Histomonas *meleagridis* is one of the most concerning disease pathogens of poultry production today, due to its severe impact on animal productivity and welfare. Since there are no approved drugs for the treatment and control of *H. meleagridis*, prevention of histomoniasis relies heavily on the control of *H. meleagridis* using anthelmintics. Consequently, failure to control FBZ-resistant *H. gallinarum* would lead to a continuous cycle of infection and disease with *H. meleagridis*. This then presents a scenario of production loss and animal welfare concerns that cannot be readily prevented.

In conclusion, we now have identified resistance to FBZ in two separate species of poultry nematodes in two successive trials. This highlights the possibility that FBZ resistance is much more common on poultry farms than is currently appreciated. Drug resistance in poultry ascarids may have important impacts both directly and, in the case of the *H. meleagridis* life cycle, indirectly on animal welfare and production loss. Given the ease with which we have found farms with drug-resistant ascarids, there is an important need to determine the scope and magnitude of this problem by investigating the prevalence of FBZ resistance in nematodes of poultry.

If the prevalence of resistance is as high as we believe it could be, it is possible that anthelminthic resistance is playing a role in the recent resurgence of *H. meleagridis* as a concern in chickens While historically not seen as a significant problem, recent evidence shows significant production impacts in broilers in layers due to histomoniasis. There are likely many complex factors contributing to this resurgence, but a lack of vector control due to anthelminthic resistance may play an important role, further highlighting the need for methods for surveillance.

Currently, we are investigating the genetic mechanisms of FBZ resistance in poultry ascarids, which appear to differ from that of strongylid nematodes. Identifying the genetic mechanisms of resistance is important, as this would facilitate the development of a diagnostic test, which would facilitate the measurement of resistance prevalence on a wide geographic scale. In addition, new alternative treatments for both for *H. gallinarum* and *H. meleagridis* are greatly needed.

## ACKNOWLEDGMENTS

We would like to think the producers and industry veterinarians for their help in identifying and procuring this isolate. We would also like to thank Pablo Jimenez Castro, Leonor Sicalo Gianechini, Kayla Dunn, and Natalie Wilson for their assistance in the study.

This project was supported by a grant from the US Poultry and Egg Association (project #F081).

## ETHICAL STATEMENT

All birds were handled under protocols approved by the University of Georgia Institutional Animal Care and Use Committee (IACUC) under animal use policy A2019 01-005-Y2-A3.

